# Reverse transcription-PCR using a Primer Set Targeting the 3D Region Detects Foot-and-Mouth Disease Virus with High Sensitivity

**DOI:** 10.1101/305086

**Authors:** Tatsuya Nishi, Toru Kanno, Nobuaki Shimada, Kazuki Morioka, Makoto Yamakawa, Katsuhiko Fukai

## Abstract

Because foot-and-mouth disease (FMD) has the potential to spread extensively, methods used for its diagnosis must be rapid and accurate. Therefore, reverse transcription-PCR (RT-PCR) plays an important diagnostic role. Here we designed the primer set FM8/9 to amplify 644 bases of the conserved 3D region of all seven serotypes of FMD virus (FMDV). We compared the performance of RT-PCR assays using FM8/9 with that using the primer set 1F/R targeting the 5’-UTR described in the manual of the World Organization for Animal Health. The detection limits of the RT-PCR assays were determined for 14 strains representing all serotypes. Compared with the sensitivities of the RT-PCR assay using 1F/R, those using FM8/9 were 10^1^-to 10^4^-fold higher for eight strains. To assess the validity of the methods for analyzing clinical samples, sera and saliva samples from pigs and cows infected with FMDV were collected daily and analyzed using the two PCR assays. The FM8/9 assay detected FMDV from all infected pigs and cows for longer times compared with the 1F/R assay, therefore revealing higher sensitivity for the clinical samples. Our results suggest that the FM8/9 RT-PCR assay is highly sensitive and is therefore suitable for the diagnosis of FMD.

## INTRODUCTION

Foot-and-mouth disease virus (FMDV) belongs to the family *Picornaviridae*, genus *Aphthovirus*. Its genome is composed of an approximately 8.4 kilobase, single-stranded, positive-sense RNA that encodes the proteins L, VP1–4, 2A, 2B, 2C, 3A, 3B, 3C, and 3D. FMDV isolates comprise immunologically distinct serotypes, as follows: O, A, C, Asia 1, and South African Territories (SAT) 1–3. Each serotype can be divided into genetically distinct topotypes and lineages (1). FMD causes the most contagious disease affecting cloven-hoofed animals, and control of FMD depends on early detection of infected animals. Therefore, rapid and accurate methods for diagnosing FMD are essential.

FMDV is detected using cell culture techniques, enzyme-linked immunosorbent assays, and nucleic acid analyses such as those described in the manual of the World Organization for Animal Health (OIE) (2). Among the latter methods, reverse transcription PCR (RT-PCR) assays rapidly detect FMDV with high sensitivity (3–5). Though the FMDV genome mutates at a high rate, similar to those of other RNA viruses (6), a highly conserved region should be selected as a target of RT-PCR to detect the seven serotypes. The OIE manual specifies a primer set for agarose gel-based RT-PCR assays (named 1F/R) that targets the 5’-UTR (7). Primer sets for real-time RT-PCR (rRT-PCR) assays that target the 5’-UTR and 3D domain are published (2, 8, 9). Although these assays rapidly detect FMDV with high sensitivity, agarose gel-based RT-PCR assays may be susceptible to cross-contamination with PCR amplicons after the reaction. In contrast, rRT-PCR tends to generate nonspecific signals, particularly from low RNA copy-number samples, and the expensive instrumentation requires strict quality control. Therefore, PCR methods should be used along with other methods to diagnose FMD (10). In the present study, the primer set FM8/9 targeting a conserved region of 3D was designed, and its performance was compared with that of the 1F/R primer set.

## MATERIALS AND METHODS

### Cells and viruses

Primary bovine kidney (BK), IBRS-2, BHK-21 and MDBK cells were grown in Eagle’s Minimum Essential Medium (Nissui Pharmaceutical, Tokyo, Japan), while ZZR-127 cells (11) were grown in Dulbecco’s modified Eagle’s medium:nutrient mixture F-12 (Life Technologies, Carlsbad, CA, USA) supplemented with fetal bovine serum. The cells were maintained at 37 °C in a 5% CO_2_ atmosphere. Virus isolation was performed according to the OIE Manual (2). The virus strain 0/JPN/2010 was isolated from clinical samples of cattle in Japan using BK cells and passed three times in BHK-21 cells. 0/M0G/4/2017 and A/TAI/46-1/2015, distributed by the State Central Veterinary Laboratory (Mongolia) and Regional Reference Laboratory for FMD in South East Asia (Thailand), were propagated using ZZR-127 cells and BHK-21 cells, respectively. Strains O/TUR/5/2009, O/ECU/4/2010, O/HKN/2015, A/TAI/2011, A/SAU/15/2016, A/IRN/1/2011, Asia1/TUR/2011, C/PHI/7/1984, SAT1/KEN/2009, SAT2/SAU/2000, and SAT3/ZIM/1983, distributed by the Pirbright institute (United Kingdom), swine vesicular disease virus (SVDV) J1’73, and two strains of vesicular stomatitis virus (VSV) of the New Jersey and Indiana 1 serotypes were grown on monolayers of IBRS-2 and/or BHK-21 cells. The bovine rhinitis virus strains A (BRAV) SD-1, M-17, and H-1 and bovine rhinitis virus B (BRBV), strain EC11, were propagated using BK and MDBK cells at 33 °C in a 5% CO_2_ atmosphere.

### RNA extraction, Reverse transcription-PCR and real-time RT-PCR

The primer set FM8/9 was designed to amplify 644 nucleotides of the 3D region conserved in all seven serotypes, based on the FMDV sequences available in GenBank (Table 1). The 271, 108, 50, 28, 14, 15, and 8 sequences of FMDV of serotype O, A, Asia 1, C, and SAT 1-3, respectively, which encompass the complete 5’-UTR and 3D regions, were downloaded from GenBank and aligned using MEGA7 software (http://www.megasoftware.net/). Nucleotide variability at each position was assessed, and the target region of FM8/9 was defined as a highly conserved region of the FMDV genome (Fig. 1).

**FIG 1.**
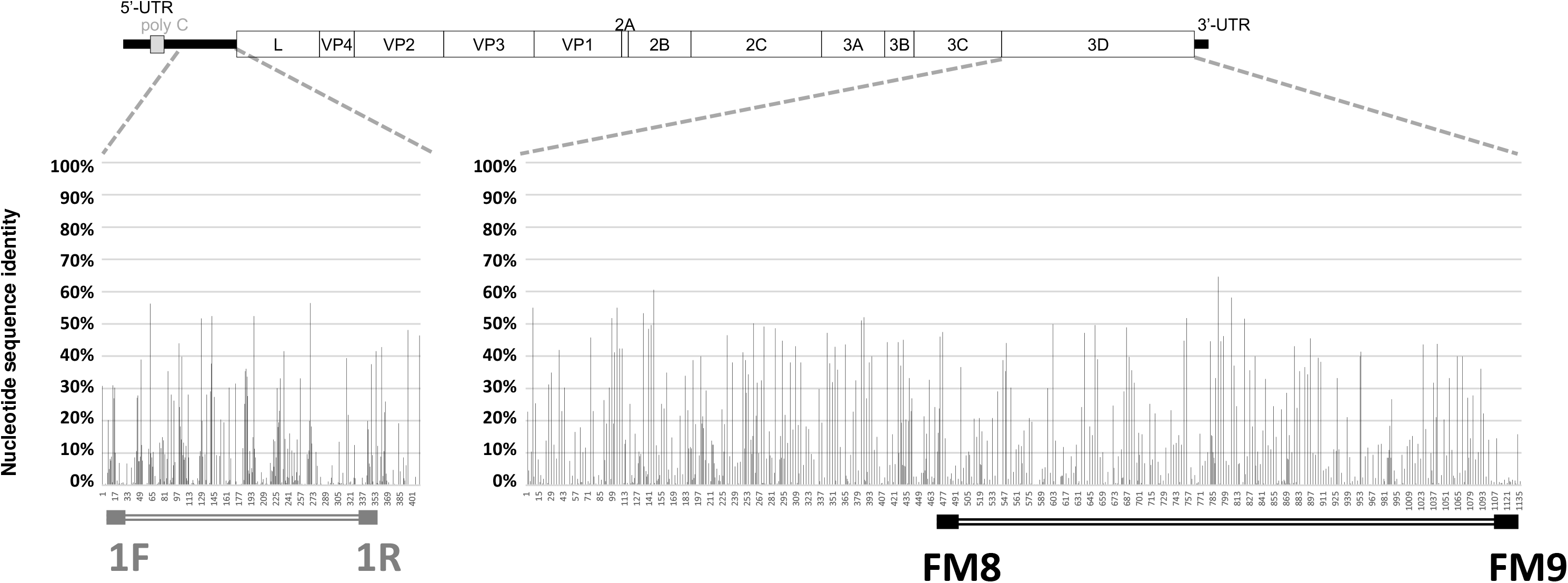
Location of RT-PCR target regions within the 5’-UTR and 3D region of FMDV. The 494 sequences of the FMDV genome, which cover the complete sequences of the 5’-UTR and 3D regions, were downloaded from GenBank. Nucleotides 278-696 of the 5’-UTR and nucleotides 1-1138 of the 3D region were compared with 0/JPN/2010-290/1E (Accession No. LC036265). The black bars show corresponding nucleotide differences. The target regions of 1F/R and FM8/9 are indicated.

**TABLE 1.**
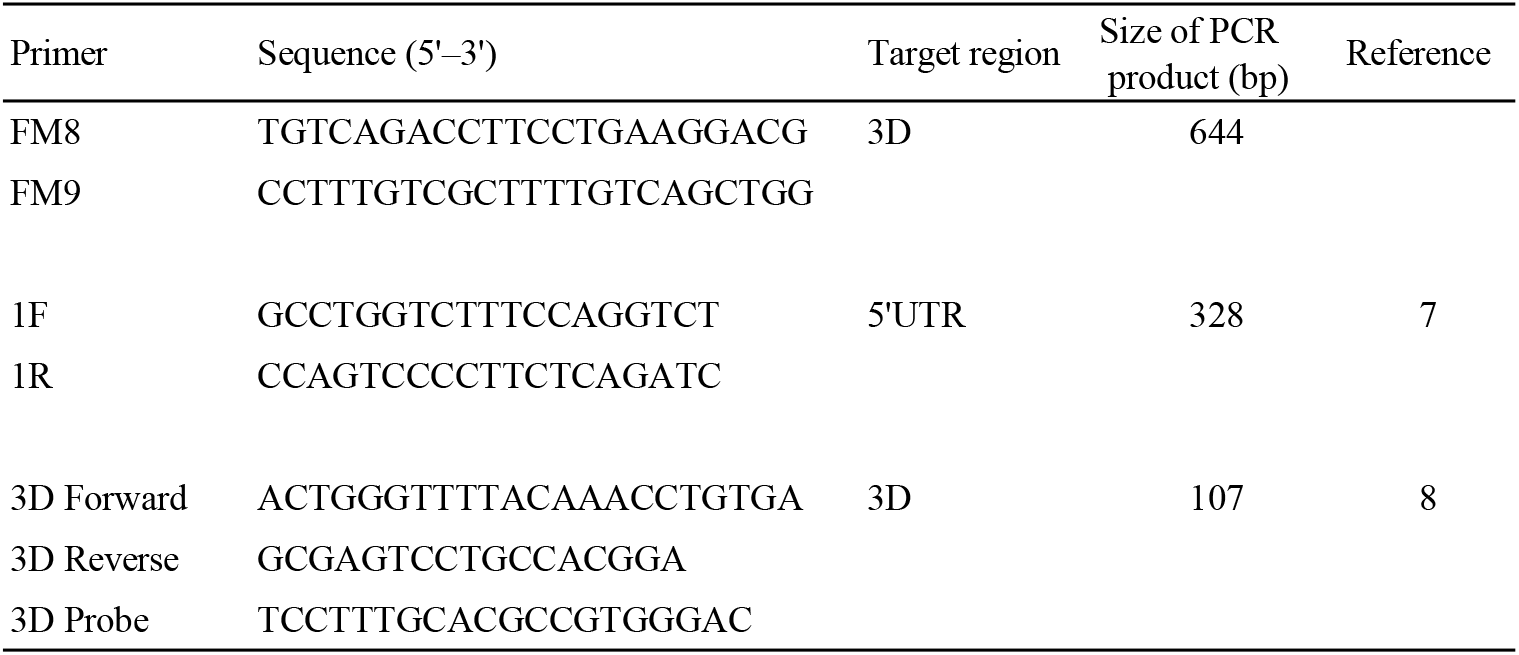
Primer sets for RT-PCR detecting FMDV genes

Viral RNA was extracted from supernatants of infected cells using the High Pure Viral RNA Kit (Roche Diagnostics, Tokyo, Japan). FMDV-specific genes were detected using a Superscript III One-Step RT-PCR System with Platinum Taq Polymerase (Life Technologies, Inc., Carlsbad, CA, USA) and primer set FM8/9 or 1F/R. PCR amplification was performed using an ABI GeneAmp PCR System 9700 (Life Technologies) as follows: 55 °C, 30 min, one cycle; 94 °C for 2 min, one cycle; 94 °C for 15 s, 55 °C for 30 s, and 68 °C for 45 s, 35 cycles; and 68 °C for 5 min, one cycle. PCR products were separated using electrophoresis through 1% agarose gels. Amplicons were stained with ethidium bromide and visualized using UV-light transillumination (Fig. 2). The rRT-PCR assay was conducted using TaqMan Fast Virus 1-Step Master Mix (Life Technologies) with the primer sets 3D-Forward, Reverse, and Probe which are described in the OIE manual (2).

**FIG 2.**
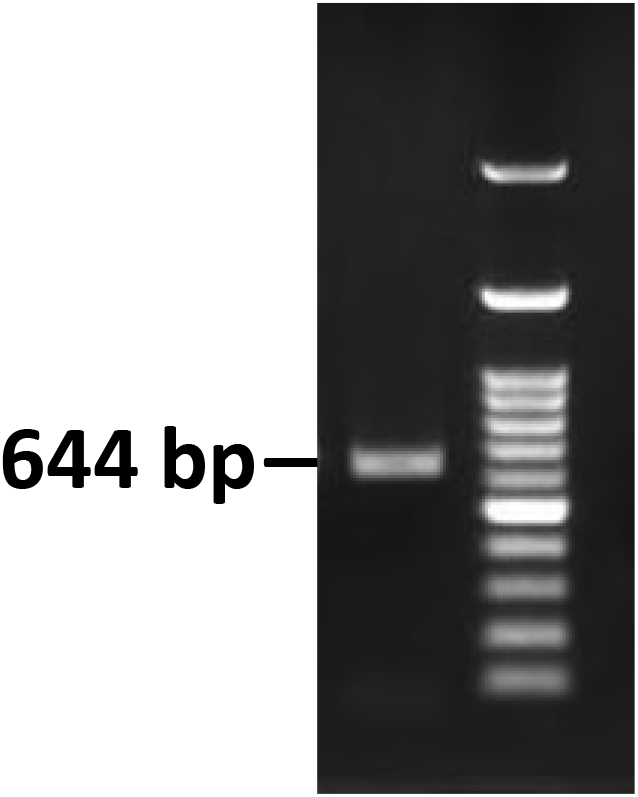
FMDV sequences amplified using RT-PCR with the FM8/9 primer set. RNA was extracted from a supernatant of cells infected with 0/JPN/2010 and subjected to RT-PCR using the FM8/9 primer set.

### Nucleotide sequencing

First-strand cDNA synthesis from the RNAs of O/JPN/2010, O/MOG/4/2017, O/ECU/4/2010, O/HKN/2015, A/TAI/46-1/2015, A/SAU/15/2016, and C/PHI/7/1984 were performed using the SuperScript III Reverse Transcriptase (Life Technologies) and FMDV-specific 2B331R primer (5’-GGCACGTGAAAGAGACTGGAGAG-3’) and 3’NT-Rc primer (5’-CAATTGGCAGAAAGACTCTGAGGCG-3’). The entire length of the L-fragment gene (approximately 7.7 kb) was amplified using PCR with PrimeSTAR Max DNA Polymerase (TaKaRa, Shiga, Japan) and two primer sets as follows: set 1, 5’-2010F primer (5’-CGTTAAAGGGAGGTAACCACAAG-3’) and 2B331R primer, and 2B217F primer (5’-ATGGCCGCTGTAGCAGCACGGTC-3’) and 3’NT-Rc primer. Double-stranded cDNAs of O/TUR/5/2009, A/TAI/2011, A/IRN/1/2011, Asia1/TUR/2011, SAT1/KEN/2009, SAT2/SAU/2000, and SAT3/ZIM/1983 were synthesized using PrimeScript Double Strand cDNA Synthesis Kit (TaKaRa). Nucleotide sequences were analyzed using the Ion PGM System (Life Technologies) as previously described (12). To analyze complete sequences of the target region of the primer set FM8/9, 3’-rapid amplification of cDNA ends (RACE) was conducted for O/ECU/4/2010 and O/HKN/2015 using a 3’-Full RACE Core Set (TaKaRa). Nucleotide sequences were analyzed using an ABI 3130 Genetic Analyzer (Life Technologies).

### Experimental infections

Two 2-month-old pigs were intradermally inoculated with 1 ml of 10^5.3^ tissue culture infectious dose (TCID_50_) of O/JPN/2010 and cohabited with four pigs (13). Two 6-month-old Holstein cows were inoculated intradermally with 1 ml of 10^7.5^ TCID_50_ of O/MOG/4/2017 and cohabited with two cows (Fukai K., unpublished observations). Sera and saliva samples were collected daily as described in our previous report (14) for 11 dpi and 10 dpi, respectively. Viral RNA was extracted from the sample as described above and analyzed using RT-PCR assays with the two primer sets. Animal experiments were authorized by the Animal Care and Use Committee of the National Institute of Animal Health (NIAH) and were performed in a high-containment facility at the NIAH.

## RESULTS AND DISCUSSION

### Detection limits of RT-PCR using primer set FM8/9 or 1F/R

We compared the detection limits of RT-PCR using the primer set FM8/9 or 1F/R to amplify the genes of 14 FMDV strains representing seven serotypes. RNAs were extracted from each serial 10-fold dilution of the 14 FMDV strains ranging from 10^−3^ to 10^4^ TCID_50_/0.1 ml titrated in IBRS-2 cell and subjected to RT-PCR and rRT-PCR (Table 2). The detection limits were decided based on the results of triplicate RT-PCR reactions. The sensitivities of the RT-PCR reaction primed by FM8/9 were >10-fold higher for detection of O/TUR/5/2009, O/MOG/4/2017, O/ECU/4/2010, A/SAU/15/2016, A/IRN/1/2011, Asia1/TUR/2011, SAT1/KEN/2009, and SAT3/ZIM/1983 (shaded gray in Table 2) compared with those using 1F/R. The detection limits for strains O/TUR/5/2009, O/MOG/4/2017, A/SAU/15/2016, A/IRN/1/2011, Asia1/TUR/2011, C/PHI/7/1984, SAT1/KEN/2009, and SAT3/ZIM/1983 were <10^−1^ TCID_50_/0.1 ml (written in bold in Table 2), indicating that the sensitivities of the RT-PCR assay were increased >10-fold compared with those of virus isolation using IBRS-2 cells. Moreover, compared with the sensitivities of the rRT-PCR assay, those of the RT-PCR assay using FM8/9 were 10-fold higher for detecting O/TUR/5/2009, Asia1/TUR/2011, and SAT3/ZIM/1983 (shaded dark gray in Table 2).

**TABLE 2.**
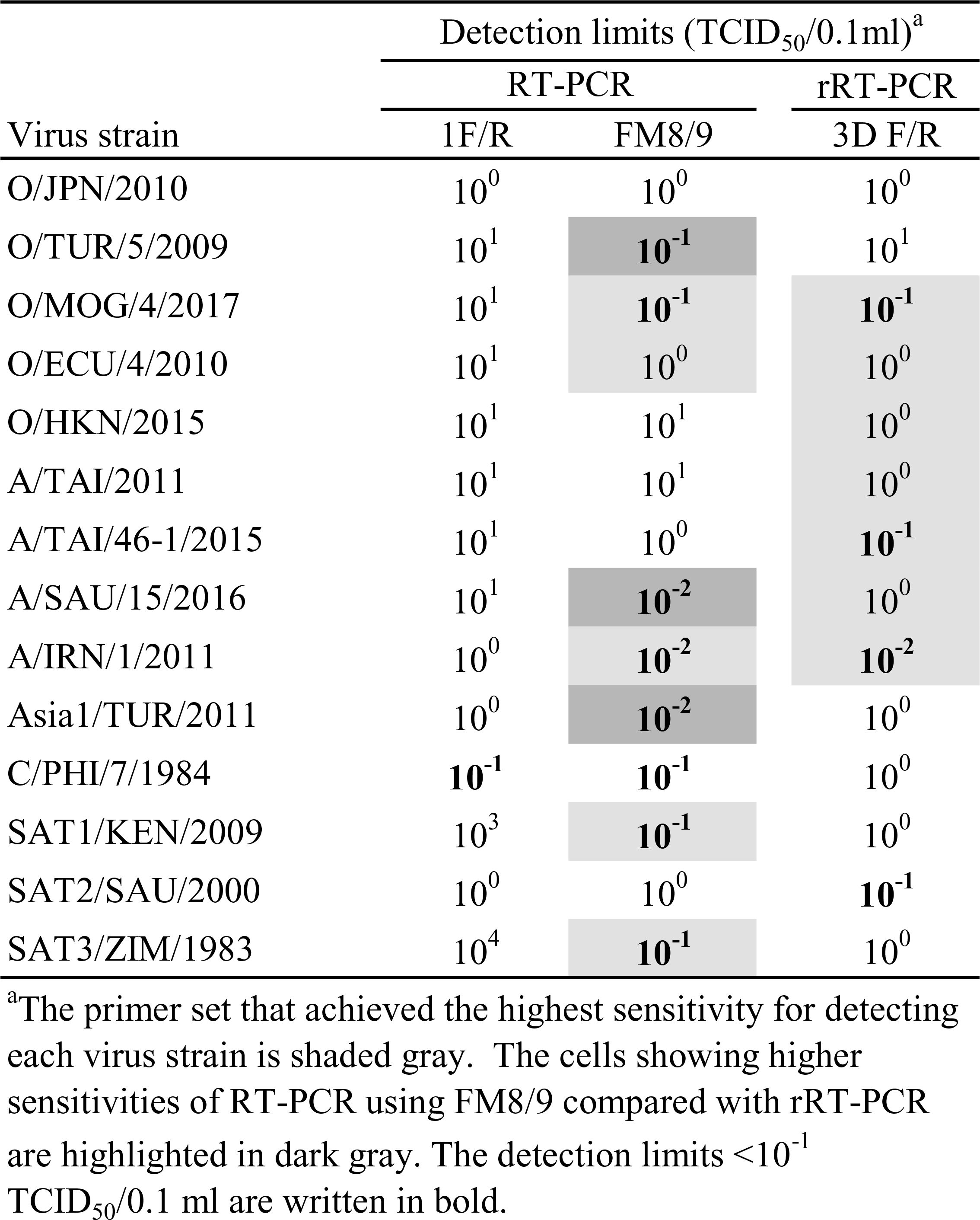
Detection limits of RT-PCR using each primer set

The nucleotide sequences of each primer’s target region of the genomes of the 14 FMDV strains are shown in Table 3. Although we assumed that the low sensitivity of the RT-PCR assay using 1F/R to detect SAT3/ZIM/1983 could be explained by two or more consecutive mismatches of the nucleotide sequences of the 1F/R primer target regions, such a significant difference in identities between the sequences of the two primer sets was not found among the other strains. These findings indicate that the sensitivities of the RT-PCR assays using each primer set may not depend on identical nucleotide sequences of a primer’s target region. Further, the Tm values and GC-contents of the primer sets, efficiencies of amplification of the target sequence, and high-order structures of RNA target region of FMDVs may influence sensitivities.

**TABLE 3.**
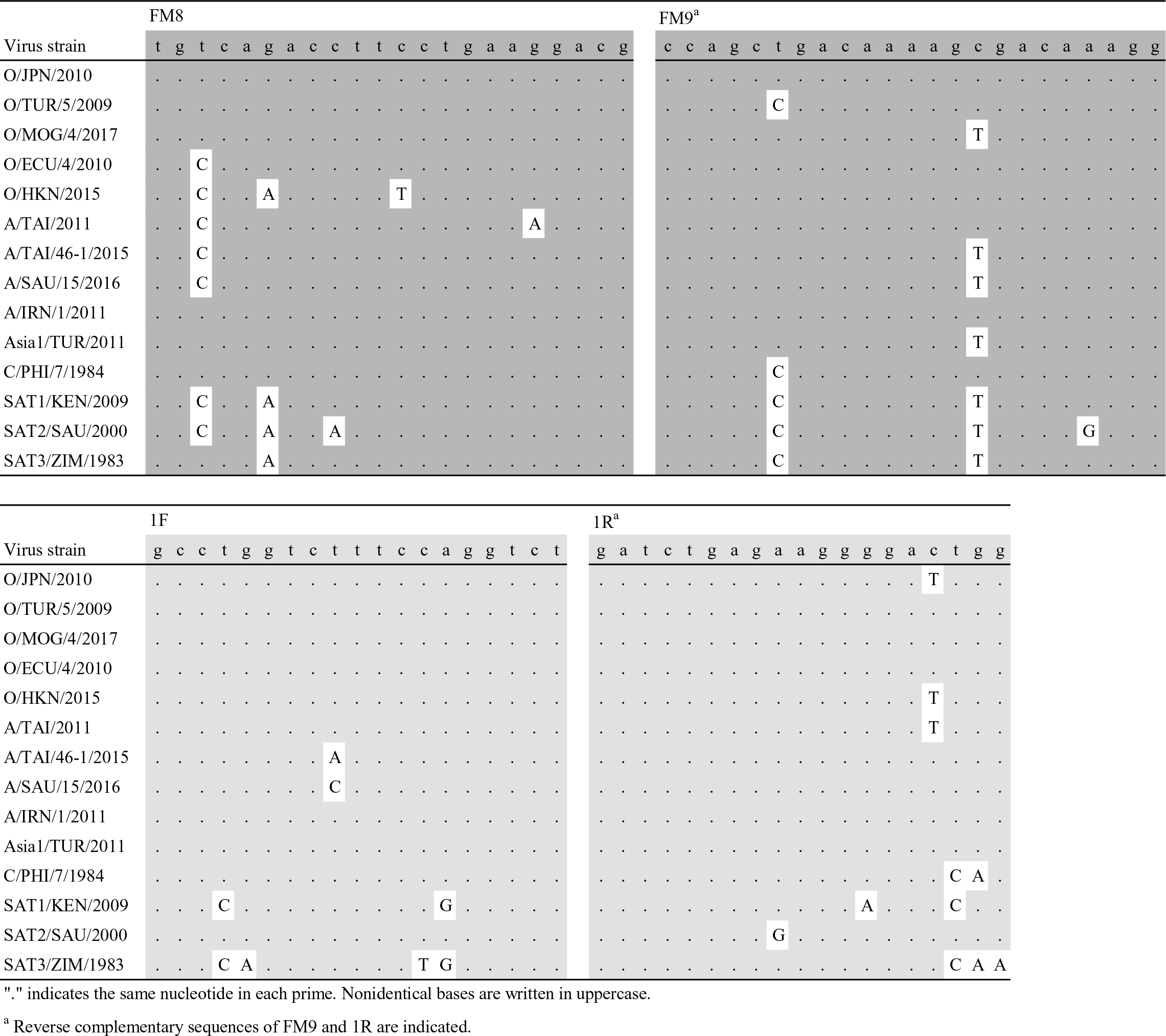
Comparison of nucleotide sequences of each primer target region of 14 FMDVs

### Validation of the specificities of primer sets for non-FMDV viruses

To validate specificity, RNA samples were extracted from SVDV strain J1’73; VSV strains New Jersey and Indiana 1 serotypes; BRAV strains SD-1, M-17, and H-1; and BRBV strain EC11. The RNAs were subjected to RT-PCR using each primer set and to rRT-PCR. There were no false-positive reactions in the RT-PCR assay using 1F/R and rRT-PCR. In contrast, the RT-PCR assay using FM8/9 yielded a weak positive reaction with BRAV strain SD-1 (data not shown). SVDV, VSV, and BRBV RNAs were not detectably amplified. Thus, diagnosis using only the RT-PCR assay with FM8/9 can lead to misidentification; however, the present RT-PCR method did not amplify BRAV genes from sera, saliva, or nasal samples 0 dpi of 32 Holstein cows (age >3 months) in the animal experiments by our institute (Table 5) (15). Further, the diagnosis of FMD in practice is determined using RT-PCR, rRT-PCR, virus isolation, nucleotide sequencing, serological tests, and epidemiological data that includes clinical signs. Clinical signs of BRV are reported to be mild respiratory disease (16, 17). Moreover, rRT-PCR, which was highly sensitive for detecting FMDV gene, did not detect BRAV, therefore, it would not be difficult to differentiate these viruses.

### Detection of FMDV genes from clinical samples using RT-PCR

To assess the validity of the RT-PCR methods for testing clinical samples, sera and saliva samples from infected and cohabited pigs and cows in previous studies of experimental infections (13, Fukai K., unpublished observations) were analyzed daily using the RT-PCR assays with the two primer sets (Tables 4 and 5). The RT-PCR assay using FM8/9 was positive from 1 dpi to 10 dpi and 1 day postcontact (dpc) to 10 dpc for inoculated and uninoculated pigs, respectively. In contrast, FMDV genes were detected from 1 dpi to 8 dpi and 2 dpc to 9 dpc using 1F/R. For samples from inoculated and uninoculated cows, the RT-PCR assay using FM8/9 detected FMDV genes from 1 dpi to 9 or 10 dpi, and from 2 dpi to 9 or 10 dpc, respectively. The RT-PCR assay using 1F/R detected FMDV genes from 1 dpi to 5 or 10 dpi, from 4 dpi to 10 dpi, or from 2 dpc to 7 dpc, respectively. These findings indicate that the RT-PCR assay using FM8/9 detected FMDV genes for a longer period from all infected pigs and cows than 1F/R. Therefore, the RT-PCR using FM8/9 achieved higher relative sensitivity for clinical samples from infected pigs and cows.

**TABLE 4.** Detection of clinical samples of FMD obtained from pigs inoculated with FMDV O/JPN/2010 and their uninoculated cohabitants

| Pig No. | Sample | Primer set | days postinoculation |  |  |  |  |  |  |  |  |  |  |  |
| --- | --- | --- | --- | --- | --- | --- | --- | --- | --- | --- | --- | --- | --- | --- |
|  |  |  | 0 | 1 | 2 | 3 | 4 | 5 | 6 | 7 | 8 | 9 | 10 | 11 |
|  |  |  |  | 0 | 1 | 2 | 3 | 4 | 5 | 6 | 7 | 8 | 9 | 10 |
| 1<br>(inoculated) | Serum | FM8/9<sup>a</sup> | - | + | + | + | - | - | - | - | - | - | - | - |
|  |  | 1F/R | - | + | + | - | - | - | - | - | - | - | - | - |
|  | Saliva | FM8/9 | - | + | + | + | + | + | + | + | + | + | + | - |
|  |  | 1F/R | - | + | + | + | + | + | + | + | + | - | - | - |
| 2<br>(inoculated) | Serum | FM8/9 | - | + | + | + | - | - | - | - | - | - | - | - |
|  |  | 1F/R | - | + | + | + | - | - | - | - | - | - | - | - |
|  | Saliva | FM8/9 | - | + | + | + | + | + | + | + | + | + | + | - |
|  |  | 1F/R | - | + | + | + | + | + | + | + | + | - | + | - |
| 3<br>(contact) | Serum | FM8/9 | NS<sup>b</sup> | - | - | + | + | + | - | - | - | - | - | - |
|  |  | 1F/R | NS | - | - | + | + | + | - | - | - | - | - | - |
|  | Saliva | FM8/9 | NS | - | + | + | + | + | + | + | + | + | + | + |
|  |  | 1F/R | NS | - | - | + | + | + | + | + | + | + | + | - |
| 4<br>(contact) | Serum | FM8/9 | NS | - | + | + | + | + | - | - | - | - | - | - |
|  |  | 1F/R | NS | - | - | + | + | + | - | - | - | - | - | - |
|  | Saliva | FM8/9 | NS | - | + | + | + | + | + | + | + | + | - | - |
|  |  | 1F/R | NS | - | - | + | + | + | - | + | + | + | - | - |
| 5<br>(contact) | Serum | FM8/9 | NS | - | - | - | - | + | + | + | + | - | - | - |
|  |  | 1F/R | NS | - | - | - | - | + | + | + | - | - | - | - |
|  | Saliva | FM8/9 | NS | - | + | + | + | + | + | + | + | + | + | - |
|  |  | 1F/R | NS | - | - | - | - | + | + | + | + | + | - | - |
| 6<br>(contact) | Serum | FM8/9 | NS | - | + | + | + | + | - | - | - | - | - | - |
|  |  | 1F/R | NS | - | + | + | + | + | - | - | - | - | - | - |
|  | Saliva | FM8/9 | NS | - | + | + | + | + | + | + | + | + | + | + |
|  |  | 1F/R | NS | - | - | + | + | + | + | + | + | + | - | - |

<sup>a</sup> Results of RT-PCR assays using FM8/9 in this table are published (13).<sup>b</sup> NS, not sampled

**TABLE 5.** Detection of FMD in clinical samples obtained from cattle inoculated with FMDV O/MOG/4/2017 and their uninoculated cohabitants

| Cow No. | Sample | Primer set | days postinoculation |  |  |  |  |  |  |  |  |  |  |
| --- | --- | --- | --- | --- | --- | --- | --- | --- | --- | --- | --- | --- | --- |
|  |  |  | 0 | 1 | 2 | 3 | 4 | 5 | 6 | 7 | 8 | 9 | 10 |
| 1<br>(inoculated) | Serum | FM8/9 | - | + | - | + | + | - | - | - | - | - | - |
|  |  | 1F/R | - | + | - | + | - | - | - | - | - | - | - |
|  | Saliva | FM8/9 | - | + | + | + | + | + | - | + | - | + | - |
|  |  | 1F/R | - | + | + | + | + | + | - | - | - | - | - |
| 2<br>(contact with<br>Cow No. 1) | Serum | FM8/9 | - | - | - | + | + | + | + | + | - | - | - |
|  |  | 1F/R | - | - | - | - | + | + | + | - | - | - | - |
|  | Saliva | FM8/9 | - | - | + | + | + | + | + | + | + | + | + |
|  |  | 1F/R | - | - | - | - | + | + | + | + | + | + | + |
| 3<br>(inoculated) | Serum | FM8/9 | - | + | + | + | + | + | - | + | - | - | - |
|  |  | 1F/R | - | + | + | + | + | - | - | - | - | - | - |
|  | Saliva | FM8/9 | - | + | + | + | + | + | + | + | + | + | + |
|  |  | 1F/R | - | + | + | + | + | + | + | + | + | + | + |
| 4<br>(contact with<br>Cow No. 2) | Serum | FM8/9 | - | - | + | + | + | + | + | + | - | - | - |
|  |  | 1F/R | - | - | + | - | + | + | + | - | - | - | - |
|  | Saliva | FM8/9 | - | - | - | + | + | + | + | + | + | + | - |
|  |  | 1F/R | - | - | - | - | + | + | + | + | - | - | - |

The present study describes a method that will likely contribute to efforts to more rapidly and accurately detect FMDV. Although the RT-PCR assay using FM8/9 amplified RNA genes from one BRAV cell culture supernatant, we demonstrated the assay’s higher sensitivities for seven serotypes of FMDVs and clinical samples from cattle and swine. In contrast, the RT-PCR using 1F/R did not detect SVDV, VSV, BRAV, or BRBV RNAs and was less sensitive for detecting FMDV RNAs. Our results suggest that the PCR assays incorporating primer set FM8/9 achieved higher sensitivity for detecting FMDV genes and are therefore suitable for the diagnosis of FMD. However, accurate diagnosis of FMD requires the use of multiple PCR methods along with other diagnostics.

## ACKNOWLEDGMENTS

We thank Yukako Hasegawa and Nobuko Saito for their technical assistance; the State Central Veterinary Laboratory (Mongolia), Regional Reference Laboratory for FMD of Southeast Asia (Thailand); and the Pirbright institute (UK) for stocks of recent epidemic FMDVs. This study was supported by a research project grant for improving food safety and animal health [Check. If this is the formal name of the grant, please capitalize the first letters.] from the Ministry of Agriculture, Forestry, and Fisheries of Japan (Project no. 2021).

## REFERENCES

1. Knowles NJ, Samuel AR. 2003. Molecular epidemiology of foot-and-mouth disease virus. Virus Res 91:65–80.

2. World Organization for Animal Health (2015) Chapter 2. 1. 5. Foot and mouth disease. Manual of Diagnostic Tests and Vaccines for Terrestrial Animals 2015. http://www.oie.int/fileadmin/Home/eng/Health_standards/tahm/2.01.05_FMD.pdf

3. Ferris NP, King DP, Reid SM, Hutchings GH, Shaw AE, Paton DJ, Goris N, Haas B, Hoffmann B, Brocchi E, Bugnetti M, Dekker A, De Clercq K. 2006. Foot-and-mouth disease virus: a first inter-laboratory comparison trial to evaluate virus isolation and RT-PCR detection methods. Vet Microbiol 117:130–40.

4. Reid SM, Ebert K, Bachanek-Bankowska K, Batten C, Sanders A, Wright C, Shaw AE, Ryan ED, Hutchings GH, Ferris NP, Paton DJ, King DP. 2009. Performance of real-time reverse transcription polymerase chain reaction for the detection of foot-and-mouth disease virus during field outbreaks in the United Kingdom in 2007. J Vet Diagn Invest 21:321–30.

5. Reid SM, Forsyth MA, Hutchings GH, Ferris NP. 1998. Comparison of reverse transcription polymerase chain reaction, enzyme linked immunosorbent assay and virus isolation for the routine diagnosis of foot-and-mouth disease. J Virol Methods 70:213–7.

6. Drake JW, Holland JJ. 1999. Mutation rates among RNA viruses. Proc Natl Acad Sci U S A 96:13910–3.

7. Reid SM, Ferris NP, Hutchings GH, Samuel AR, Knowles NJ. 2000. Primary diagnosis of foot-and-mouth disease by reverse transcription polymerase chain reaction. J Virol Methods 89:167–76.

8. Callahan JD, Brown F, Osorio FA, Sur JH, Kramer E, Long GW, Lubroth J, Ellis SJ, Shoulars KS, Gaffney KL, Rock DL, Nelson WM. 2002. Use of a portable real-time reverse transcriptase-polymerase chain reaction assay for rapid detection of foot-and-mouth disease virus. J Am Vet Med Assoc 220:1636–42.

9. Reid SM, Ferris NP, Hutchings GH, Zhang Z, Belsham GJ, Alexandersen S. 2001. Diagnosis of foot-and-mouth disease by real-time fluorogenic PCR assay. Vet Rec 149:621–3.

10. King DP, Ferris NP, Shaw AE, Reid SM, Hutchings GH, Giuffre AC, Robida JM, Callahan JD, Nelson WM, Beckham TR. 2006. Detection of foot-and-mouth disease virus: comparative diagnostic sensitivity of two independent real-time reverse transcription-polymerase chain reaction assays. J Vet Diagn Invest 18:93–7.

11. Brehm KE, Ferris NP, Lenk M, Riebe R, Haas B. 2009. Highly sensitive fetal goat tongue cell line for detection and isolation of foot-and-mouth disease virus. J Clin Microbiol 47:3156–60.

12. Nishi T, Yamada M, Fukai K, Shimada N, Morioka K, Yoshida K, Sakamoto K, Kanno T, Yamakawa M. 2017. Genome variability of foot-and-mouth disease virus during the short period of the 2010 epidemic in Japan. Vet Microbiol 199:62–67.

13. Fukai K, Morioka K, Yoshida K. 2011. An experimental infection in pigs using a foot-and-mouth disease virus isolated from the 2010 epidemic in Japan. J Vet Med Sci 73:1207–10.

14. Fukai K, Yamada M, Morioka K, Ohashi S, Yoshida K, Kitano R, Yamazoe R, Kanno T. 2015. Dose-dependent responses of pigs infected with foot-and-mouth disease virus O/JPN/2010 by the intranasal and intraoral routes. Arch Virol 160:129–39.

15. Onozato H, Fukai K, Kitano R, Yamazoe R, Morioka K, Yamada M, Ohashi S, Yoshida K, Kanno T. 2014. Experimental infection of cattle and goats with a foot-and-mouth disease virus isolate from the 2010 epidemic in Japan. Arch Virol 159:2901–8.

16. Mohanty SB, Lillie MG, Albert TF, Sass B. 1969. Experimental exposure of calves to a bovine rhinovirus. Am J Vet Res 30:1105–11.

17. Betts AO, Edington N, Jennings AR, Reed SE. 1971. Studies on a rhinovirus (EC11) derived from a calf. II. Disease in calves. J Comp Pathol 81:41–8.

